# Looming detection in complex dynamic visual scenes by interneuronal coordination of motion and feature pathways

**DOI:** 10.1101/2023.09.20.558565

**Authors:** Bo Gu, Jianfeng Feng, Zhuoyi Song

## Abstract

Detecting looming signals for collision avoidance faces challenges in real-world scenarios due to interference from moving backgrounds. Astonishingly, animals, like insects with limited neural systems, adeptly respond to looming stimuli while moving at high speeds. Existing insect-inspired looming detection models integrate either motion-pathway or feature-pathway signals, remaining susceptible to dynamic visual scene interference. We propose that coordinating interneuron signals from the two pathways could elevate looming detection performance in dynamic conditions. We used artificial neural network (ANN) to build a combined-pathway model based on *Drosophila* anatomy. The model exhibits convergent neural dynamics with biological counterparts after training. In particular, a multiplicative interneuron operation enhances looming signal patterns. It reduces background interferences, boosting looming detection accuracy and enabling earlier warnings across various scenarios, such as 2D animated scenes, AirSim 3D environments, and real-world situations. Our work presents testable biological hypotheses and a promising bio-inspired solution for looming detection in dynamic visual environments.

---

Detecting looming signals, the expanding moving patterns generated by incoming objects, lies at the heart of collision avoidance. Yet, real-world challenges arise as the agent’s motion generates dynamic background movements, bewildering the foreground object features and corrupting those vital looming signals (Fig. 1). Remarkably, animals, like insects with simple brains, excel at responding to looming stimuli while moving swiftly. Behavioural experiments show that even flying flies, the simplest insect with only 100,000 neurones in the brain, can react to looming stimuli with ultra-fast visually directed banked turns^1^.

**Fig. 1.**
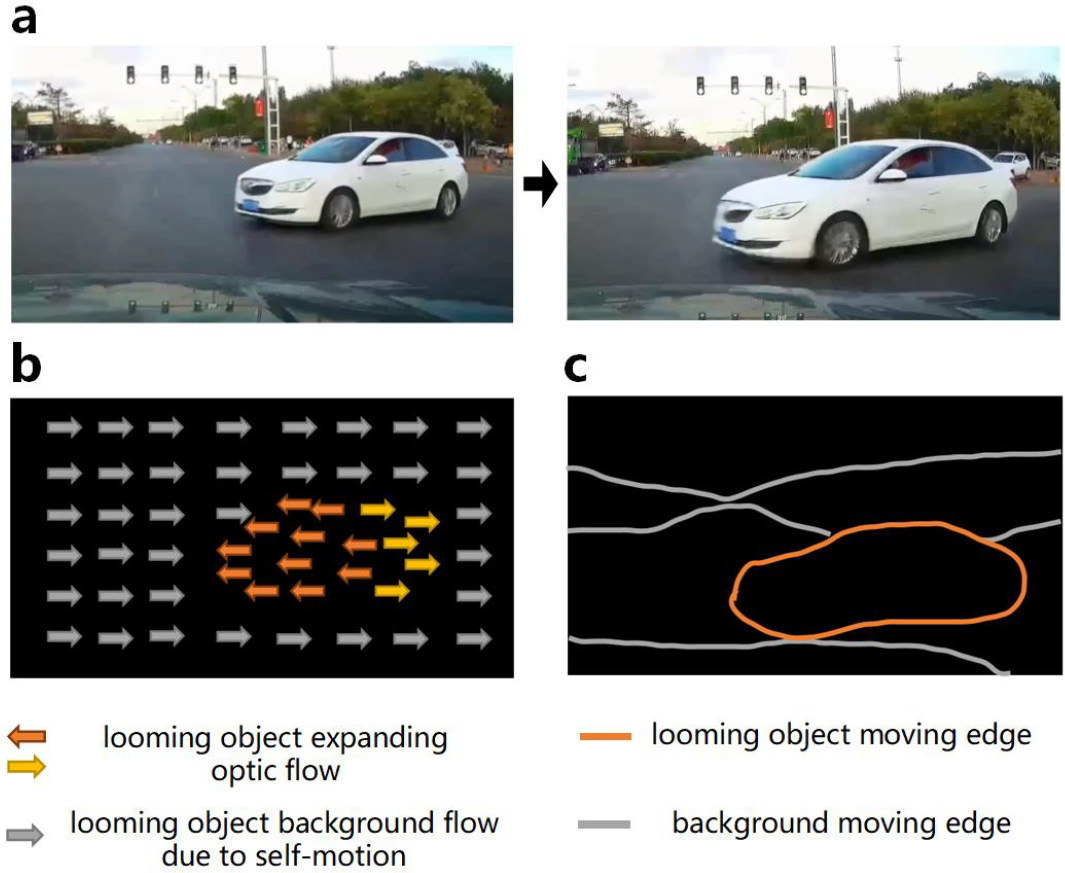
Dynamic backgrounds create interference in looming estimation. (a) A car approaching in a natural scene; (b) The optic flow of sequential images shows the car’s optic flow mixed with background optic flow, making it challenging to distinguish the car’s looming pattern; (c) The moving edges in the visual space are unclear, as the car’s edges blend with the environment’s edges.

In the spirit of this extraordinary natural talent, the realm of neuro-inspired computations^2,3^ has found innovative applications in artificial systems such as automobiles and unmanned aerial vehicles (UAVs) to address challenges of collision avoidance^4,5^. This integration has led to significant reductions in computation time while showcasing impressive achievements in detecting imminent obstacles^6^. However, the pursuit of perfection remains ongoing.

Although traditional solutions excel in uncomplicated settings or static natural scenes^7^, they stumble when confronted with complexity - especially scenarios where foreground objects and backgrounds dynamically intertwine^8^. In this particular investigation, we introduce an innovative neuroinspired computational framework driven by recent biological insights. This framework is poised to revolutionise the detection of looming signals in visually dynamic environments.

The core of a looming signal lies in how its subtended angle on the observer (*θ*) changes over time^9^. Thus, looming detection can be categorised into two theoretical schemes: one based on calculating changing image features and the other on detecting motion patterns. Similarly, insect-inspired looming detection models fall into two types: motion-pathway models and feature-pathway models, depending on whether they integrate local motion signals or contrast edge changes to form a looming selective response. This classification traced the evolution of insect-inspired looming detection models over the past 30 years.

The first insect-looming detector, called the LGMD model^10,11^, is a feature-pathway model named after the lobula giant movement detector (LGMD) neurone in the locust visual system. It is a shallow neural network with only four to five layers, using local illumination changes as inputs rather than explicitly coding directional motion signals^12^. The key to its looming detection lies in the excitation-inhibition network structure. By striking a delicate balance between excitation and delayed inhibition, the model selectively responds to expanding objects while ignoring receding ones. This selectivity is crucial for detecting specific features.

The LGMD model relies on critical image cues for detecting impending objects, such as speed increases of an edge motion or size increases of an expanding edge. However, a challenge with edge detection is its vulnerability to irrelevant background and object features, leading to false alarms for nearby or accelerating translating stimuli^3^. Researchers have tried various enhancements to improve the accuracy and robustness of LGMD models in complex visual scenes. For example, some attempted to group nearby excitations^2^, while others explored ON/OFF pathway competitions^**?**,13^, added a separate contrast pathway to scale the looming responses^14^, or introduced competitions between local excitation and inhibition signals^15^. Despite these efforts, background feature interference continues to be a persistent issue in these models.

An alternative network structure was proposed two decades ago, integrating local directional motion signals for looming detection^16^, even without biological evidence at the time. Subsequent research validated the importance of local motion detectors in *Drosophila* for computing expansion flow fields^17^. More recent experimental findings^18^ and models^**?**,14^ have provided insights into the workings of the looming detector neuron, LPLC2 (lobula plate/lobula columnar, type II), in *Drosophila*. This neuron integrates local motion signals and selectively responds to the diverging expanding motion pattern for effective looming detection. However, this model structure prioritises ultra-selective responses to the radial motion expansion when an object approaches head-on, potentially compromising robustness in detecting various looming patterns from different directions.

Could the moving-edge feature detection pathway and the motion-dependent flow-field calculation coordinate for looming detection? How do they coordinate? Are there extra functional benefits for looming detection in dynamic backgrounds? This paper investigates these questions by constructing an artificial neural network model (ANN) constrained by *Drosophila* neuroanatomy. This ANN aims to detect looming stimuli in dynamic visual scenes.

Drawing insights from novel connectomic data in *Drosophila*^19^, the looming detection circuits receive synaptic inputs from both the motion and feature detection pathways. A novel multiplication operation is introduced between the interneuron signals from these pathways, creating a combined-pathway model. The ANN encompasses the flexibility of reducing to either a motion-pathway or a feature-pathway model if looming detection exclusively utilises motion or feature detection inputs without multiplication. Moreover, to mimic the remarkable abilities of various neuron types in processing distinct motion patterns that coexist in the visual scene, the ANN includes parallel branches to handle wide-field background motion, translating object motion, and looming motion patterns, respectively. This analogy culminates in the ANN to mirror the multifaceted dynamics of real-world visual scenes.

To optimize performance, the ANN undergoes supervised training with a multi-objective function, which weights the detection of various motion patterns. Finally, the trained ANNs are evaluated in different conditions, including looming in 2D animated static and dynamic backgrounds, AirSim simulated 3D environments and real-world situations. The performance of the combined-pathway model is compared to the other two models, the motion-pathway and feature-pathway models.

Astoundingly, the ANN converges to solutions exhibiting similar neural dynamics as real biological counterparts after training. A captivating revelation unfolds in the results: while the integration of two pathways yields negligible benefits and seems superfluous in simple or static backgrounds, it becomes an exhilarating catalyst in dynamic realms. The combined-pathway model’s accuracy in looming signal estimation soars by up to 30 percent in dynamic backgrounds. Furthermore, the combined-pathway model bestows the gift of much earlier warnings in both simulated and real-world scenarios. This extraordinary performance boost stems primarily from the ingenious multiplication operation. It empowers the model to magnify the looming signal pattern while deftly suppressing interference noise generated by unpredictable background movements. This study showcases the pivotal role of interneuronal coordination in motion and feature detection signals, serving as a potent catalyst for augmenting looming detection efficacy in intricate dynamic scenes.

## Related work

Insect’s visual system is a refined biological structure that can efficiently implement survival tasks. *Drosophila* is a traditional model organism with a tiny brain but complicated behaviours, contributing to the science community with excellent experimental tools in genetics, anatomy and physiology. More importantly, for computational purposes, stereotyped neural circuits come in handy for mechanistic modelling. Here, we briefly review related looming detection models and neural circuits, focussing on the implementations with complementary pathways.

### Complementary motion and feature pathways in *Drosophila* neural circuit

In *Drosophila*, visual information is processed through four layers of neuropils in its optic lobe before being sent to the central brain and integrated with information from other sensory modalities^**?**,**?**,20^. These four optic lobe neuropils are the retina, lamina, medulla, and lobula complex. The lobula complex is further divided into two distinct structural loci: the lobula plate and the lobula providing parallel motion and feature processing pathways that coordinate behaviours.

The neurones in the lobula plate specifically code motion signals, such as optical flow information in the visual field^21^. There are two parallel motion pathways that exist for ON and OFF motion, emerging as early as in the Lamina, but showing motion-sensitive responses in T4 and T5 neurones within the lobula plate. T4 and T5 cells integrate excitation or inhibition inputs from adjacent column signals to form direction selectivity, coding local motion signals^22^. Furthermore, they send their dendrites to four separate layers in the lobula plate, coding moving signals in four coordinated directions, respectively^12^. Thousands of T4 and T5 cells tile the whole visual space, forming an optic flow map, like ganglion cells in vertebrates. The lobula plate tangential cells (LPTC) then integrate these local motion signals to create a sense of the global pattern induced by self-motion.

On the other hand, the lobula is a feature-detecting structure responsible for recognising behaviourally relevant visual objects, such as conspecifics or predators^**?**^. Visual projection neurons in the lobula, like lobula columnar (LC) and lobula tangential (LT) cells^23^, send signals to specific brain regions coding various visual features^24^. For example, LC18, LC21, and LC11 can detect small moving objects that extend 2-4 degrees and even code the object’s moving speed with linear speed tuning curves^25^.

Although the lobula and the lobula plate are densely connected, indicating some interdependence between parallel processing of motion and feature, complementary signals may assist in certain behavioural functions. For example, looming detection circuitry involving LPLC1 and LPLC2^23^ may show this cooperation. LPLC2 selectively integrates local motion signals from T4 and T5 neurones to detect radial expanding motion patterns, indicating an approaching object on a collision course^18^. LPLC1, however, receives input from both motion pathways and feature detection interneurons (T2 and T3), triggering slowing behaviours upon stimuli of back-to-front motion, mimicking a frontal or parallel approaching object^26^. Unlike well-studied T4 and T5 neurones, T2 and T3 cells are only gaining more research attention these years.

T2 and T3 neurons have tight size tuning curves and provide excitatory presynaptic inputs to object-selective visual projection neurons^**?**,27^. They were previously modelled to protrude the outline of a small object within a small moving target detector (STMD) circuit^24,27^. Their integration with T4 and T5 cells in the looming detection circuit and the extra functional benefits they provide are still not fully understood. We hypothesise in this paper that combining T2&T3 and T4&T5 signals may enhance looming detection performances in dynamic visual scenes.

### Insect biology-inspired movement detector models

Researchers have created numerous bio-inspired mechanistic models for movement detection in insects. However, most of these models perform well only in simple laboratory settings due to difficulties in distinguishing object features from complex backgrounds. To address this, continuous efforts have been made to enhance the models’ performance by introducing new specific mechanisms or pathways. Here, we briefly highlight the improvements made in three classes of related models: elementary motion detectors, looming detectors, and small target moving detectors.

### Wide-field movement detectors

Lobula plate tangential cells (LPTCs) in the fly’s visual system process wide-field motion signals. They integrate local motion inputs coded by T4 and T5 cells, which can be modelled as elementary motion detectors (EMDs). Research suggests that LPTCs have receptive fields similar to matched filters for various optical flow patterns^28^. Although theories on LPTC computations have not seen much progress since the early 2000s, discoveries in neurological implementations of EMD circuits have advanced rapidly. These include the ON-OFF pathway split^**?**^, separate four layers of T4 and T5 cells for different coordinates^12^, and hybrid motion computations involving prefered direction enhancement and non-prefered direction inhibition^**?**,29^. These new models enhance direction selectivity and reduce output ambiguities, leading to improved optical flow estimations^30^.

### Looming detectors

Insect looming detectors were established in the mid-1990s and have seen significant advancements in recent years. There are two types of looming detectors, LGMD models for the locust visual system^10,11^ and LPLC2 models for *Drosophila*^**?**,14^. The key difference between them is whether local motion signals are explicitly coded in the network.

Both the LGMD and LPLC2 models face challenges from background interference in dynamic visual scenes. Researchers have proposed combining feature-based and motion-based models to improve the performance of various motion patterns. Yue proposed two decades ago to integrate a translating-sensitive neural network (TSNN) to extract visual motion cues from the whole field^**?**^, even though they came to the conclusion that such a redundant scheme does not add much to the LGMD performance^**?**^. Q.Fu integrated an LPTC model^**?**^ and an LGMD model to enhance the model’s looming detection selectivities, reducing false alarms to nearby translating stimuli^**?**^. More recently, Shuang et al. combined the two types of looming detection models to achieve both image velocity selectivities of LGMD and the radial motion opponency characteristics of LPLC2^8^.

However, most of these previous attempts integrated different models independently and introduced competition mechanisms in the output. Instead, we believe that a more effective approach is to introduce interaction among the pathways at the level of interneurons, creating a more reliable and efficient neural circuit.

### Small target movement detectors

Our model also takes inspiration from the Small Target Moving Detection (STMD) model, which is a crucial set of object detection models^31,32^. Although not typically discussed with LGMD or LPLC models, recent studies in *Drosophila* suggest correlations between them. Neurons in the lobula, like LC18 and LC11, have been identified as STMDs. LC11 and LPLC1 may even share interneurons that encode the outline features of an object^19,27^. The classic STMD model detects fast-moving objects by leading and trailing edges that have opposite directional contrast changes^33^. Modifications have been made to account for new experimental findings, such as LC18’s smaller tuning size for small targets, which was addressed by introducing a bounded crossover inhibition mechanism^24^. On the other hand, the selectivities for small targets of LC11 neurones were successfully replicated by introducing spatial-temporal pooling^19^. These novel STMD models rely on T2 and T3 to provide the moving edge signal before the correlation step^27^. We speculate that moving edges may also enhance looming object detection in dynamic backgrounds in *Drosophila*.

### Biological anatomy-constrained neural networks

The mechanistic models discussed earlier are carefully designed to replicate the responses of specific neurons, but they may only work well under specific conditions. On the other hand, Artificial Neural Networks (ANNs) trained on datasets can perform efficiently with numerous parameters but lack interpretability. Recently, neural scientists have started constraining ANNs with real neural circuit anatomy^30,34,35^, showing that such models can converge to biological solutions and be more efficient and robust^**?**^. This convergence of biological neural circuits and artificial neural networks can mutually validate their rationality under realistic conditions.

In the context of movement detectors in *Drosophila* neural circuits, several anatomy-constrained ANNs have been developed. For example, Mano et al. created a shallow convolutional ANN that models the processing steps of the elementary motion detection circuit^30^. Their model was trained in moving scenes to estimate movement speed using T4 and T5 signals. Drews et al. developed a similar ANN based on T4 and T5 neurons with divisive normalization, showing that spatial contrast normalization improves motion estimation on different natural scenes^36^. Zhou et al. established a two-layer convolutional ANN for the looming-detection neuron LPLC2, successfully distinguishing between looming objects that will hit or miss^26^. However, there has yet to be an ANN modelling the elementary motion detection, looming, and translating object detection circuits together to investigate their interaction effects.

## Results

### Anatomically constrained ANN model for looming detection in dynamic visual scenes

In real-world dynamic scenes, various motion patterns and the *Drosophila* neural circuits have evolved parallel branches to handle them. Our model also incorporates parallel branches for wide-field background motion, translating object motion, and looming motion patterns (Fig. 2a). These branches share the same interneurons (T2, T3, T4, and T5), making the network compact.

**Fig. 2.**
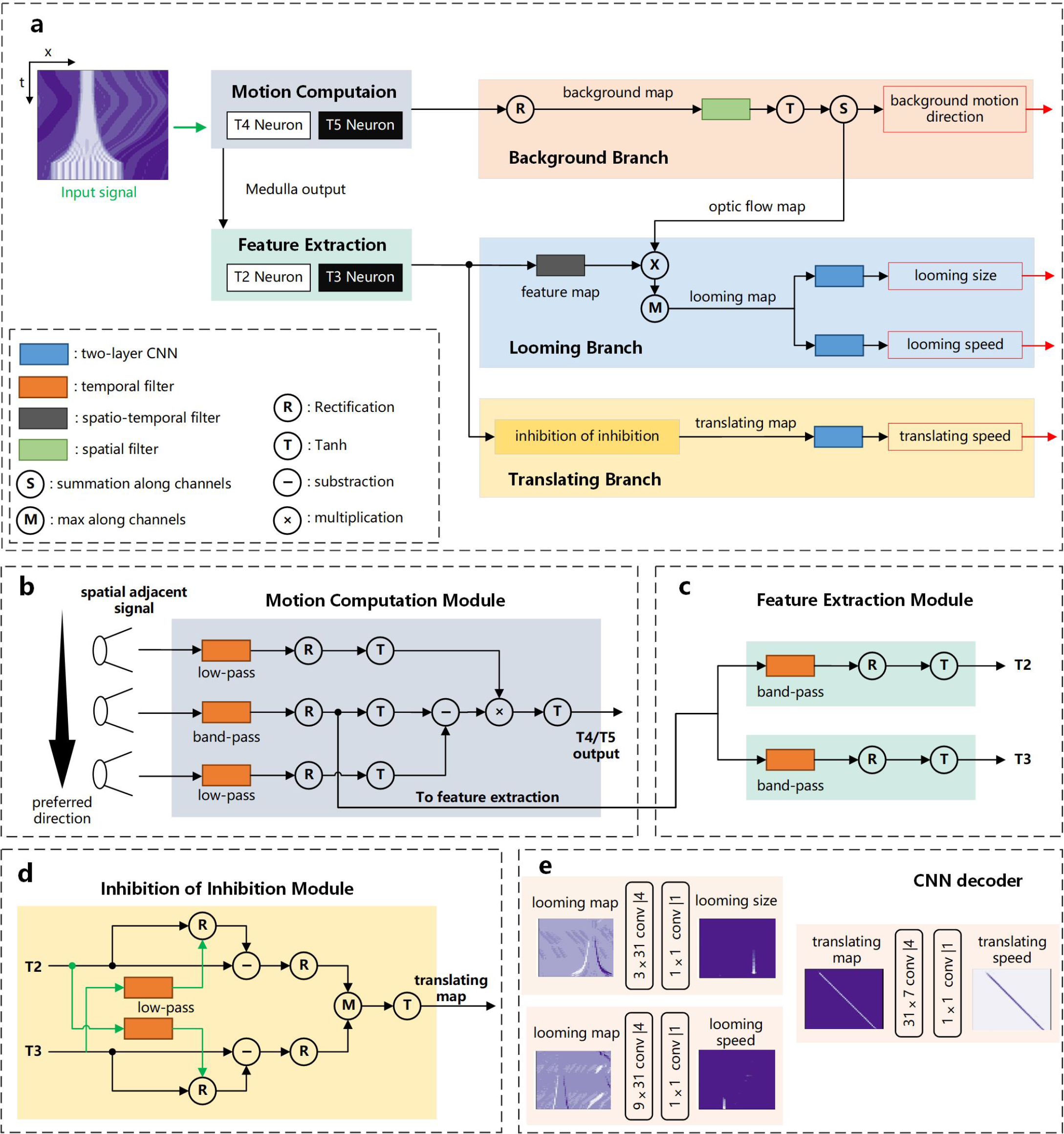
Model Framework. (a) The model consists of three branches for background motion, translating object speed, and looming objects’ size and speed detection, respectively. The looming branch combines signals from motion-detection interneurons (T4 and T5 cells) and feature-detection interneurons (T2 and T3 cells). (b) The elementary motion detection model calculates the local motion direction by integrating three adjacent column signals. (c) The inhibition of inhibition module is based on the LC18 neuron model. This module computes the trailing edge of the translating object using ON and OFF moving edge signals.

The background branch estimates the direction of the background motion using local motion signals from T4 and T5 neurones^30^. We used a basic EMD model^29^ with static compression for this branch (Fig. 2b). The translating branch takes inputs from T2 and T3 neurones and uses an STMD structure to detect small translating objects (Fig. 2c,d). The STMD model adopts ideas from the inhibition of inhibition mechanism from neural models for LC18 in *Drosophila*^24^. We then calculate the translating object’s speed using a two-layer CNN. The looming detection branch combines input from all four types of interneurons to acquire a looming map that shapes the expanding pattern of the looming object. On the basis of this map, we compute the object’s expanding size and speed^37^ using two-layered CNNs (Fig. 2e).

The interactions between T2 / T3 and T4 / T5 in *Drosophila* are not fully understood. We speculate that they sharpen each other’s signals and filter out noise. Thus, we introduce a multiplication operation to combine the visual information from T2/T3 and T4/T5. This makes the ANN a combined-pathway model with multiplication, but it can be reduced to a motion-pathway or feature-pathway model if only T4/T5 or T2/T3 signals are used for looming detection. We train the spatio-temporal receptive fields of all interneurons using a loss function that considers errors from all three branches. This multi-objective optimisation allows the network to converge to biologically plausible solutions and alleviates the fine-tuning problem when each branch is trained separately.

#### Visual stimuli dataset for training the ANN model

Our model was trained and tested on videos that contain both foreground moving objects and background moving scenes (Extended Data Fig. 1). We used panoramic high-dynamic range images as the background and manually added looming or translating foreground objects. For simplicity and generality, we focused on the 1D scenario, using a line of pixels as our raw data (Extended Data Fig. 1a left). The motion of this pixel line created a 2D spatial-temporal pattern, which was labelled with background motion directions, looming size, looming speed, and translating speed. In total, we generated 2886 training samples and 972 testing samples.

To test the robustness of the looming detection models in various dynamic scenes, we varied the statistical properties of both the background and foreground objects. We used 1) static backgrounds as a control and dynamic backgrounds that can move with different speeds, including stochastic speeds or uniform speeds at 0.5, 1, or 1.5 times the linearly expanding looming speed. Detecting a looming object becomes more problematic when the background moves at a comparable speed. 2) The foreground translating objects had randomly generated sizes (20-35°) and were scattered in random positions, moving leftward or rightward against the background. 3) Looming objects approached with either uniform or decreasing speeds, resulting in either nonlinear expanding or linear expanding looming areas on the retina (Extended Data Fig. 1b). Exponentially expanding looming signals are harder to detect early. 4) Looming objects had either uniform or sinewave pattern contrast; patterned looming objects were more challenging to detect than uniform objects. 5) We carefully controlled the object’s contrast to mimic predators in nature, where predators from bright skies or dark caves may have distinct contrasts against the background.

#### Interneuron responses to looming and translating objects show expected neural dynamics

We first investigated the interneuron responses of our ANN models to confirm their similarity to biological counterparts (Extended Data Fig. 3). We tested the models with four types of moving objects in the visual field: ON and OFF looming objects, and ON and OFF translating objects. The four types of interneurons encode distinct visual features. T2 and T3 mainly focused on the object’s moving edge, responding little to the background motion (T2 and T3 in Extended Data Fig. 3d) except when a high-contrast object was present (T2 and T3 in Extended Data Fig. 3c). T2 and T3 separated into ON/OFF pathways like T4 and T5, meaning they responded to specific contrasted looming objects or edges of translating objects. T2 detected the ON leading edge of an ON translating object (T2 in Extended Data Fig. 3c) or the ON trailing edge of an OFF translating object (T2 in Extended Data Fig. 3d), while T3 responded to the OFF leading edge of an OFF translating object (T3 in Extended Data Fig. 3d) or the ON trailing edge of an ON translating object (T3 in Extended Data Fig. 3c). When a looming object was present, T2 and T3 detected the expanding pattern for ON and OFF looming objects (T2 & T3 in Extended Data Fig. 3a&b), respectively.

On the other hand, T4 and T5 responded to local motion signals. They were each sensitive to ON or OFF looming objects and showed little response to objects with non-preferred contrast. T4 responded to an ON looming object with a clear looming pattern (T4 in Extended Data Fig. 3a), while T5 coded OFF looming objects (T5 in Extended Data Fig. 3b). They both exhibited motion direction opponency, i.e. positive and negative responses for the two edges that move in opposite directions, illustrated by the bright and dark strips for leftward and rightward moving edges in their feature map. For translating objects, T4 responded to the ON leading edge of an ON-contrast object or the ON trailing edge of an OFF-contrast object (T4 in Extended Data Fig. 3c&d), while T5 responded reverse (T5 in Extended Data Fig. 3c&d). Compared to T2 and T3, T4 and T5’s responses were more crowded due to background optical flow. The functions of these interneurons were biologically interpretable and aligned with our expectations.

#### The combined-pathway model demonstrates significant looming estimation enhancement and allows earlier warnings in dynamic backgrounds

We then compared the performance of three models: combined-pathway, motion-pathway, and feature-pathway models, in estimating looming objects on testing data. The combined-pathway model demonstrated significantly improved performance regardless of whether the looming object approached with a linear (Extended Data Fig. 4) or nonlinear speed (Fig. 3). All three models performed well on static background datasets, suggesting that discriminating foreground motion against static backgrounds is not particularly challenging for the models (the zero columns in Fig. 3b-d).

**Fig. 3.**
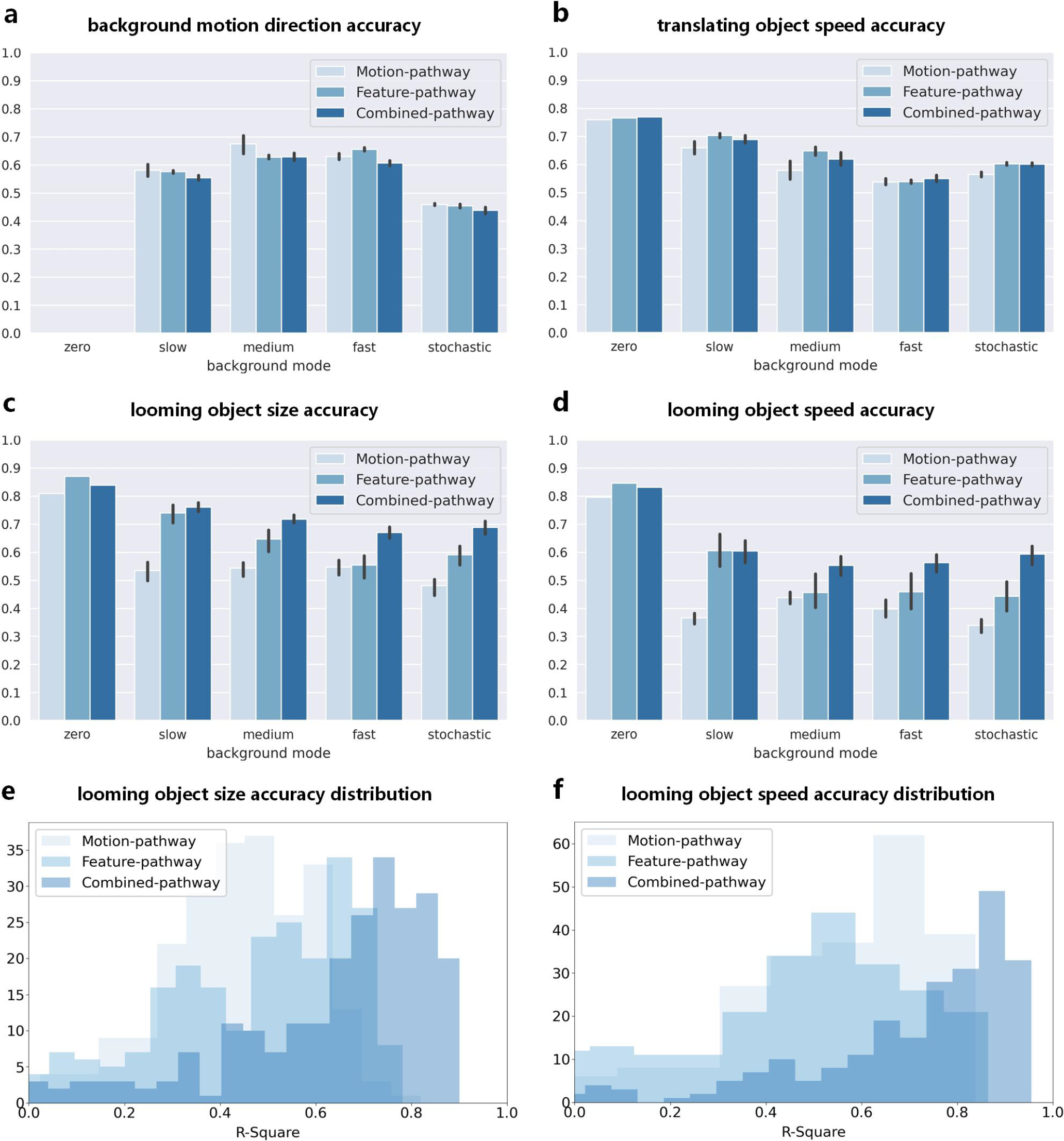
Comparisons of three looming detection model performances in static or dynamic backgrounds with exponential expanding objects, moving stochastically or with uniform speeds at different ratios of the linearly expanding looming speed. All three models’ performances decline significantly when the background is moving. However, the combined-pathway model enhances looming detection performances in all tested dynamic conditions. (a b) The three models show similar performance for estimating background motion direction and translating object speed, indicating that the complementary mechanism does not negatively affect these two branches’ functions. (c d) The complementary model improves the accuracy of estimating looming object size and speed in all tested dynamic conditions but not in static backgrounds. (e f) Performance distributions for looming size and speed estimation for the three models tested with stochastic moving backgrounds.

However, dynamic backgrounds posed challenges for the models in various motion detection tasks. Background direction estimation became harder with more dynamic background movements, especially with stochastic moving backgrounds (Fig. 3a). Moreover, all three models showed decreased performance in estimating foreground object motions when backgrounds were moving (Fig. 3b-d). Looming detection tasks were significantly more demanding in dynamic backgrounds than static backgrounds, with the motion-pathway model’s looming speed estimation accuracy dropping to only about 30% to 40% (Fig. 3d). The feature-pathway model generally performed better than the motion-pathway model, likely because of the salience feature of looming objects. Interestingly, in all looming detection conditions tested with dynamic backgrounds, the combined-pathway model outperformed the other two models, achieving a 10-25% increase in estimation accuracies (Fig. 3c-d). Particularly with faster background movements, the combined-pathway model’s accuracy in speed estimation improved by at least 50 percent compared with the other two models (Fig. 3d).

Of course, the models’ performances varied when evaluated with different image statistics, which caused different background interferences. As shown in the varied performance distributions for looming speed estimation in stochastic moving backgrounds (Fig. 3e&f), the estimation accuracy distribution of each model is approximately unimodal with slight asymmetries, certifying the broad image statistics covered. In agreement with training performance, the combined-pathway model had the highest R-square score, followed by the feature-pathway model and then the motion-pathway model. Moreover, the combined-pathway model exhibited a more constrained distribution variance, making it more robust across varied background statistics.

Lastly, detecting collisions or predators in the early stage is essential for survival or flight safety. The combined-pathway model saw looming objects earlier than the other two models, which is another significant improvement. When stimulated with an ON or OFF looming object, the combined-pathway model produced temporal responses that closely matched the ground truth in all cases (responses selected around the looming object’s central location in Fig. 4). In contrast, the motion-pathway and feature-pathway models were not sensitive in the early stage of the looming object’s approach, showing a delay in response until the object’s size reached 75° in visual space. This delay might be because the object’s small size made its edges difficult to distinguish from the background flows. The combined-pathway model overcame this challenge and provided responses when the looming object was still small.

**Fig. 4.**
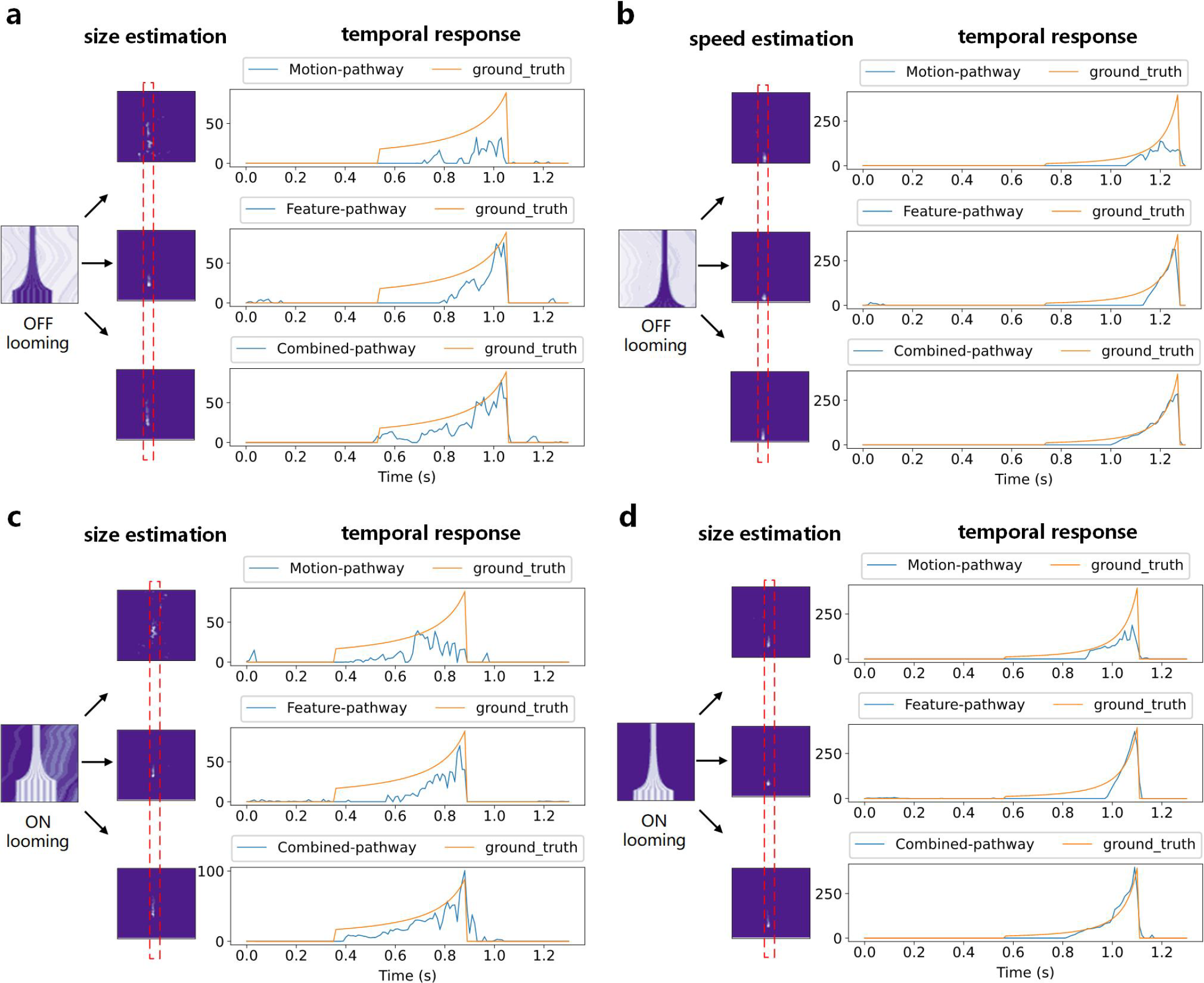
The combined-pathway model excels in early estimation of a looming object’s size and speed. The three models were tested on a specific data sample, selected to match the R-score difference at their peak distributions of the motion-pathway and combined-pathway models. (a) Size estimation map and temporal responses for an ON looming object; (b) Speed estimation map and temporal responses for an ON looming object; (c) Size estimation map and temporal responses for an OFF looming object; (d) Speed estimation map and temporal responses for an OFF looming object.

#### The multiplication of interneuronal signals is the key for enhanced looming estimation performance

We investigated why the combined-pathway model outperforms in looming estimation. Observing the results in Fig. 3, it became evident that the model’s improvement lies in the post-processing of interneuron signals, i.e. T4, T5, T2 and T3 signals. All three models have similar performances for estimating the background motion direction and the translating object speed (Fig. 3a-b), indicating that the complementary mechanism does not reversely impact the training of the background and translating object branches. This means that the combined-pathway model should have converged similar interneuron functions, compared with them trained independently on separate tasks.

To validate this, we compared the interneuron responses and the looming object feature map. We used tSNE mapping to visualize the interneuron responses from six independently trained models for each model type. Four distinct clusters emerged, corresponding to the four interneuron types, while data points for different model types were mixed (Fig. 5a). In contrast, the tSNE mapping of the looming object feature map showed different model types clustering together (Fig. 5b). This converse cluster feature illustrated that the multiplication operation between the feature map of T2&T3 signals and the optical flow map of T4&T5 signals boosted the looming estimation performance of the combined-pathway model.

**Fig. 5.**
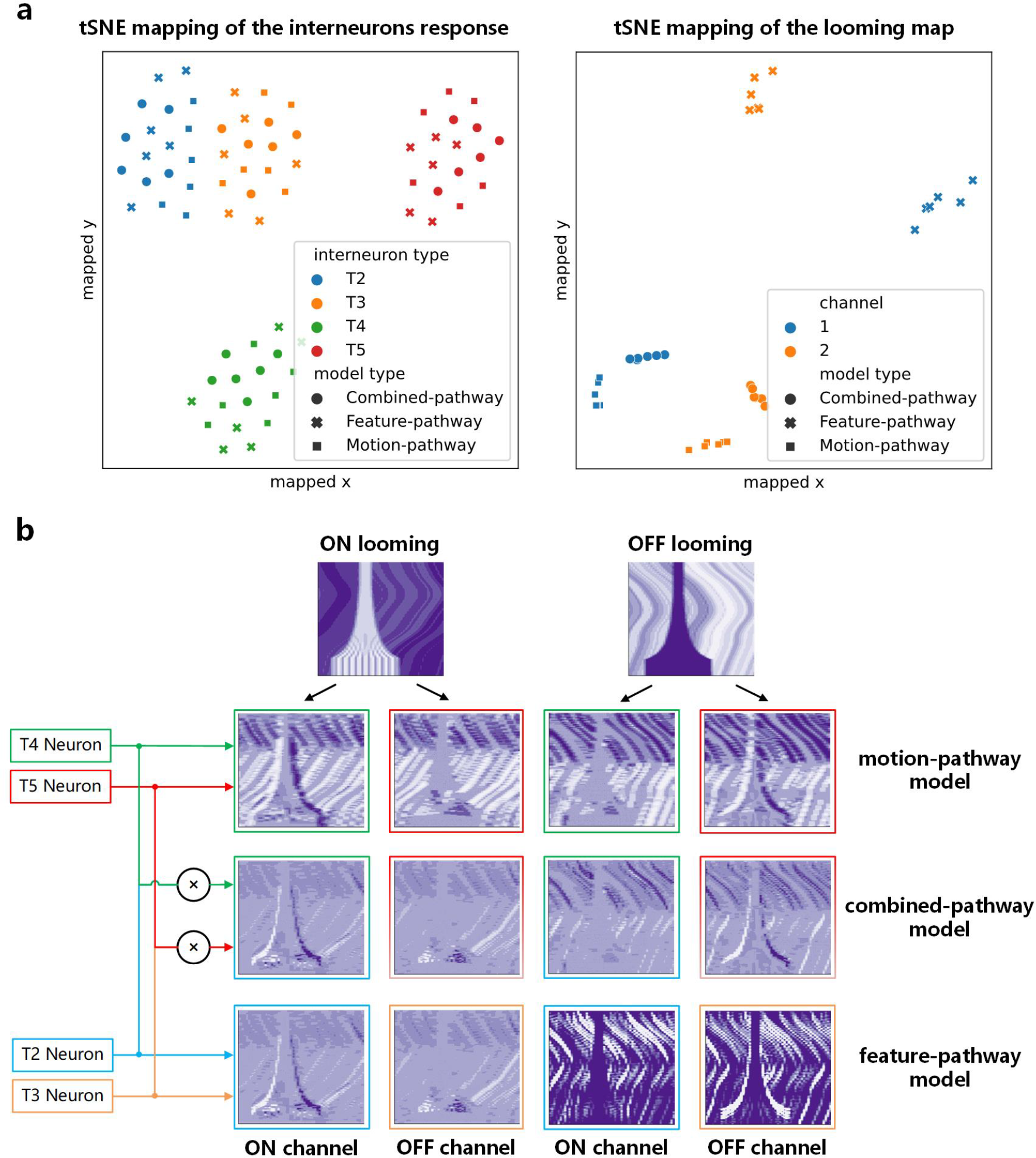
The multiplication is the key to enhancing looming estimation performance in the combined-pathway model. (a) Four clusters for the four types of interneurons show in the tSNE mapping of interneuron signals from the three model types, whereas data points for different model types are muddled together; each model type contains six independently trained models. (b) Three clusters for the three model types show in the tSNE mapping of the looming feature map. (a) and (b) indicates that multiplication is why the combined-pathway model has enhanced looming detection performance. (c) The multiplication enhances the looming signal pattern while reducing interference noise from background movements. The motion-pathway model’s looming object feature map is crowded with background optic flows. The feature-based model’s looming object feature map is cleaner but doesn’t have direction opponency. The complementary model integrates the advantages of the other two models, inducing a less noisy and direction-opponent looming object feature map.

The multiplication operation proved useful because it enhanced the selectivity for the looming object while reducing interference noise from background movements (Fig. 5b). The motion-pathway model, which only used T4 and T5 signals, suffered from a crowded looming object map due to background flows, although direction opponency helped improve selectivity. The feature-based model, relying on T2 and T3 signals, had a cleaner looming object map but lacked motion direction selectivity, leading to confusion with high-contrast background objects and reduced robustness against dynamic backgrounds. Additionally, the feature-pathway model’s responses were prominent only when the looming object was close, causing small responses in the early looming phase when the object was still small. In contrast, the multiplication allowed the combined-pathway model to inherit advantages and avoid drawbacks from the other two models. The looming signal pattern in the optical flow and feature maps showed significant correlation, and the multiplication enhanced the correlated looming signal while retaining motion direction opponency. Conversely, background motions and features produced sparse complementary interferences, which were reduced through multiplication.

#### Testing with AirSim simulated 3D scenes and real-world looming senarios

Considering that all three models were trained on synthetic images with simple contrasted blocks imitating looming objects, we asked how well our results could be generalised to more realistic scenarios. We tested the three trained models on AirSim simulated 3D scenes^38^ and real-world looming data recorded with a cell phone. In the AirSim simulated data, a flying agent moves in a selected 3D environment, while the installed camera records the scene from its own perspective. We then added an expanding contrast block with a sinewave pattern, simulating the approaching looming object. To imitate the different looming patterns caused by frontal or sideway approaching objects, we make the fly move in two distinct ways in the 3D scene. The agent can either move forwardly, with a looming object coming in a direct collision course or move left, with a looming object coming sideways (Fig. 6a). In real-world scenes, a person carries a cell phone to record the scene while he/she is walking towards a particular object. As we recorded the videos in daylight, there was a dark looming object simulating the flight course toward a collision.

**Fig. 6.**
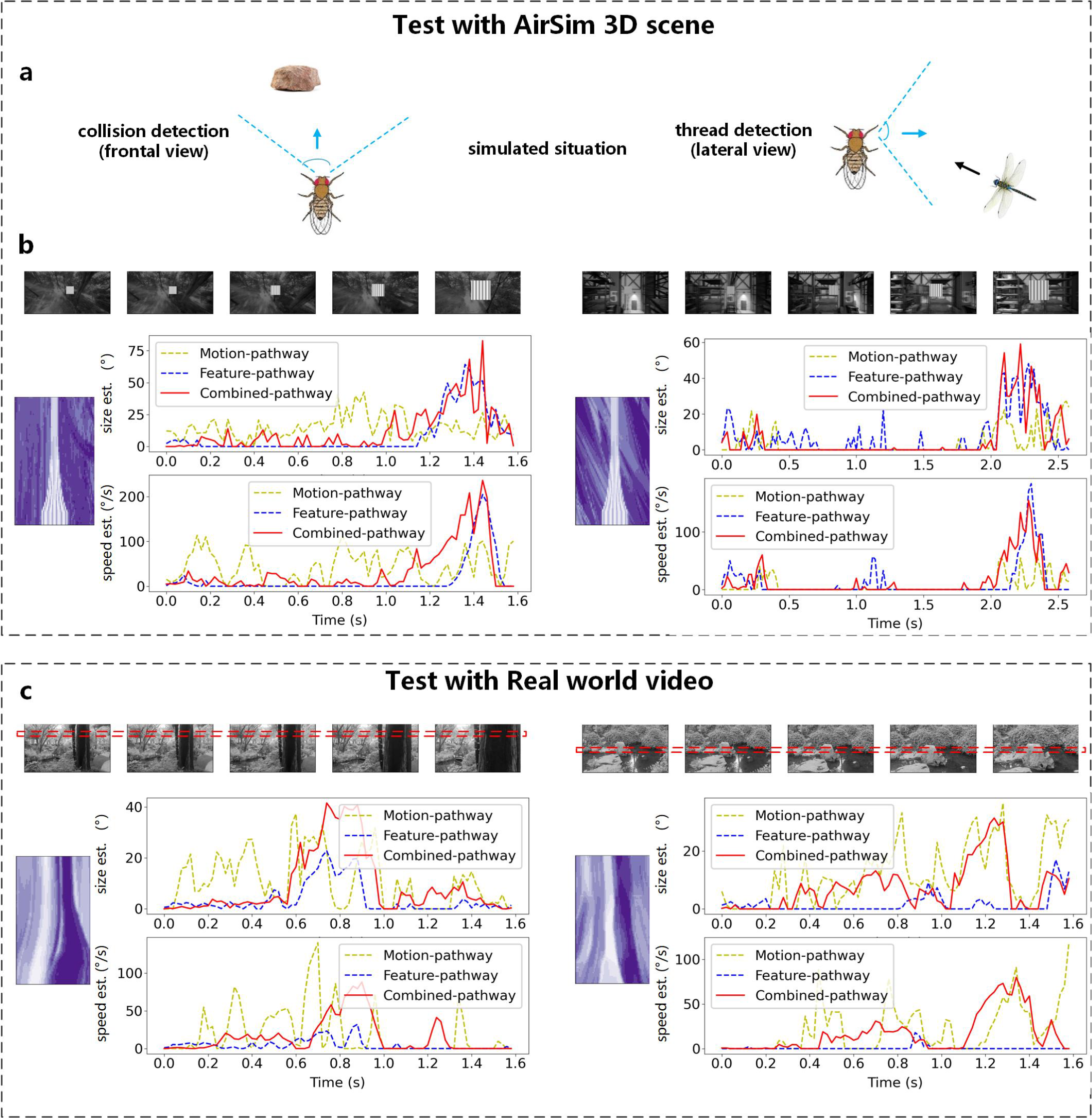
The complementary model outperforms in looming estimation in both AirSim simulated 3D dynamic visual scenes and real-world scenarios. (a) Linearly expanded looming object estimation in AirSim simulated scenes; (b) Exponentially expanded looming object estimation in AirSim simulated scenes; (c-f) Videos recorded with a cell phone in the real world with a dark looming object.

The AirSim simulated data differ from the previous synthetic training data in the more realistic background movements and the more diversified moving objects’ velocities caused by various depths. Accordingly, the looming responses become much noisier (Fig. 6b&c) compared with that tested with 2D image movements, where the looming estimation curves show rather clear exponential rising trends with little noise (ground truth in Fig. 4). This comparison highlights that movements embodied within a 3D scene can cause more background interference to looming signals than pure 2D image movements. Interestingly, background interference affected the models differently depending on whether the looming was frontal or sideways (Fig. 6b&c). The motion-pathway model was noisier during the frontal approach (Fig. 6b), while the feature-pathway model had more noise during the sideways approach (Fig. 6c). One can not simulate the converse effects with simple 2D image movements, where the motion-pathway model’s responses are always noisier (Fig. 3). This difference also reinforces the importance of testing in an embodied 3D environment.

Despite the challenges, the combined-pathway model still outperformed the others, predicting looming size and speed more accurately, reducing noise, and allowing earlier detections. For frontal approach looming signals, the combined-pathway model produces much less noise than the motion-pathway model, and it can predict the looming signal earlier in its coming than the feature-pathway model (Fig. 6b). Similarly, for side-way approaching looming signals, the combined-pathway model produces less noise than the feature-pathway model, and it can boost the faint looming signals more than the motion-pathway model (Fig. 6c). Surprisingly, the complementary mechanism’s benefits were more evident when the looming object moved slowly in its early approaching phase. When the object moved fast in the last stages, the feature-pathway model performed comparably well in estimating the looming size and speed (blue lines in Fig. 6b). However, at slower speeds, the motion-pathway model produced little noise but failed to detect looming (yellow lines in Fig. 6b), while the feature-pathway model had a larger response but was extremely noisy (blue lines in Fig. 6c), leading to potential false alarms. The combined-pathway model enhanced the looming response and suppressed noise, compensating for the limitations of each pathway (blue lines in Fig. 6b&c).

Real-world scenes presented even greater challenges due to complexities like uneven contrast edges and off-centre looming objects with edges expanding at inhomogeneous speeds. Thus, the temporal characteristics of looming estimation became more complicated. Despite this, the combined-pathway model continued to outperform the others, offering clearer speed estimations and reduced noise (red lines in Fig. 6d&e).

Our model’s performance in both AirSim simulated data and real-world looming recordings demonstrated its adaptability to complex scenarios. These tests certify the general benefits of looming estimation by coordinating signals from both the motion and the feature pathways. Note that the models were trained on synthetic datasets without retraining on new data; they could potentially perform even better if trained on more realistic datasets in the future, though this was beyond the scope of our current research.

## Discussion

Detecting impending collisions amidst dynamic real-world scenarios remains a formidable challenge, complicated by the ever-changing complexities of moving backgrounds that obscure looming objects. Yet, nature’s marvels never cease to amaze as even tiny-brained insects respond swiftly to looming threats during high-speed flights. In pursuing bioinspired solutions, existing looming detection models rooted in motion or feature pathways still contend with interference in dynamic scenes.

This study unveils a promising approach to elevate looming detection in dynamic backgrounds by harmoniously coordinating feature and motion detection pathways. Drawing inspiration from recent *Drosophila* neuroanatomy, we crafted a neural network model and honed its skills through various motion detection tasks. By creatively accepting synaptic input from motion and feature detection neurones, the looming detection branch embraces the power of pathway coordination. A seemingly simple yet potent mathematical operation — the multiplication of interneuron signals — unleashes an elegant prowess, gracefully empowering the model to counter background disturbances. This synergy culminates in heightened accuracy for looming estimations and early collision warnings amidst real-world scenes adorned with realistic looming objects.

Moreover, the model’s triumph is further enriched by a parallel structure, adeptly handling diverse motion patterns, including wide-field background motion, translating object motion, and looming motion patterns. Notably, the optimization process, fueled by multiple objective functions, deftly considers various motion patterns simultaneously. This emphasizes the significance of designing training tasks for bio-inspired neural networks targeted to solve real-world problems. Overall, this proposed looming detection model offers a promising bioinspired solution for collision detection in the real world, while simultaneously suggesting testable hypotheses in biological experiments.

## Methods

### The background branch and the EMD

The background branch (blue branch in Fig. 2a) processes local motion signals from T4 and T5 cells using neuroinspired mathematical operations to estimate the direction of the whole field motion, which has a ground truth label. The difference between the estimation and the ground truth contributes to a multi-objective loss function that tunes the parameters for interneuron filtering profiles in the motion computation module.

The input is a preprocessed high-resolution image contrast signal *I*_*c*_(*x, t*). To mimic the low resolution of the *Drosophila* compound eye, *I*_*c*_(*x, t*) was convolved with a Gaussian-shaped receptive field whose FWHM and intervals are both 3 degrees^20^:

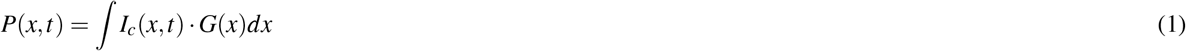

where *P*(*x, t*) denotes the optic signals received by each ommatidium in the compound eye. We integrated this step in the process of generating the dataset rather than in the runtime, saving much redundant computation during training.

Following the columnized structure in *Drosophila’s* optic lobe, each pixel corresponds to a “column” structure in the Laminar, where computations are mostly independent across columns (Fig. 2b). Here, signals are filtered and rectified into light increments and decrements^39^, representing the well-known ON/OFF pathway split in the laminar. The computations on the ON and OFF pathways are independent. Mechanisms may be implemented differently in biology; we used identical mathematical operations for these two pathways for simplicities.

The signals from adjacent laminar columns are then filtered in the Meddula and integrated to estimate the local motion directions in the lobula plate. We used an EMD model with three input arms: leading, central, and trailing edges^**?**^, based on recent neuroscience findings. These arms have varied temporal dynamics^29^, which delay and filter the signals to enhance or separate spatial-temporal correlations caused by motion. Typically, the leading and trailing edge signals are much delayed compared to the central ones:

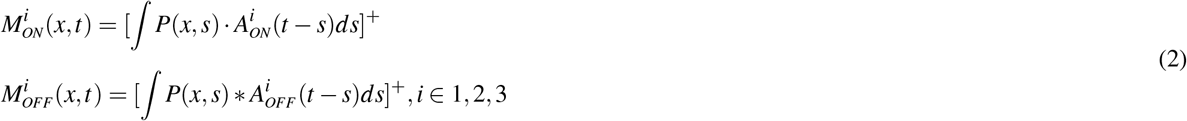

where *M*_*ON*_ and *M*_*OFF*_ are the ON/OFF pathways output signals. 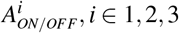 are the Medulla temporal filters for the three adjacent arms in ON/OFF pathways. [ ]^+^ represents the rectification process. For each of these arms, we also add a Tanh nonlinearity to account for the contrast adaptation in Medulla.

In our approach, we use the EMD model structure, but instead of manually setting up the medulla filters, we train them to optimize the task. Previous studies have shown that training the medulla filters can lead to biologically meaningful results, even when guided by neural signals from only one T4 or T5 neuron^30^. In the final step of EMD, the signals from the three arms are combined to enhance the preferred direction and inhibit the nonpreferred direction along the local movement direction. Traditionally, this is achieved using a correlation or multiplication between the trailing and central edges, similar to the Hassenstein-Reichardt Correlator (HRC) model, followed by a division between the central and leading edges, similar to the Barlow-Levick model. However, recent research suggests that other nonlinear signal combinations can also work well for the three-input model^40^. To avoid training difficulties associated with division operations, we used subtraction to model the inhibition instead.

In Fig. 2b, we show the basic idea for one-dimensional motion detection, and we can formally express the leftward and rightward motion outputs as follows:

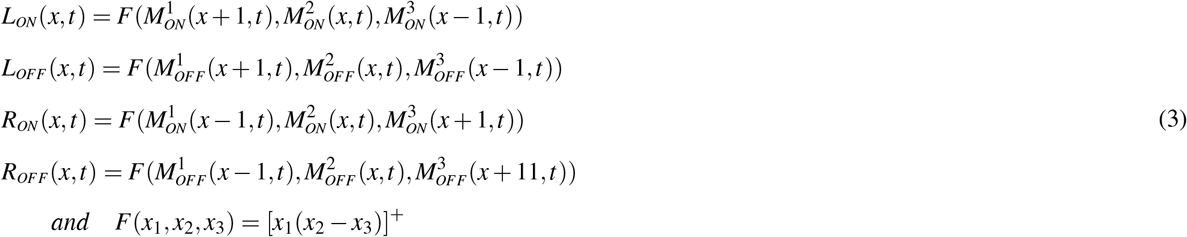

where *F* is the spatial correlation function, *R*_*ON/OFF*_, *L*_*ON/OFF*_ is the rightward or leftward motion output from ON/OFF pathways.

After spatial correlation, the signals undergo another Tanh non-linearity to obtain a normalised direction estimation. To filter out noise, a spatial filter that covers three columns is applied. Finally, we subtract the leftward and rightward outputs to obtain a signed value for direction estimation:

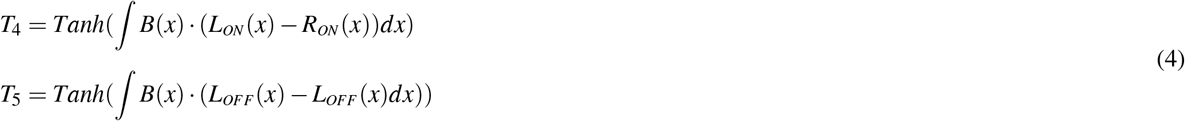

where *B* stands for the spatial filter, and *Tanh* stands for the tanh-function nonlinearity.

#### The translating object detection branch and the moving edge sensitive neurons

We added a new branch called the translating object detection branch, which helps train the feature-detection interneurons’ filtering profiles and assists in looming object detection (yellow branch in Fig. 2a). To estimate the moving speed of a translating object, we use a two-layer CNN that takes the translating object map as input. The translating object map is a feature map that marks potential objects.

To calculate the translating object map, we hypothesized that T2 and T3 neurons in the Locula play a role (Fig. 2c). These neurons are sensitive to local high-contrast changes caused by moving objects, assisting small object feature detection. Moreover, they are not direction-selective, suggesting they are feature-detection neurones. Additionally, T2 and T3 neurons provide inputs to looming detection neurons like LPLC1.

We took inspiration from the STMD model for *Drosophila* LC18 neuron, a small moving object detection neuron that receives inputs from T2 and T3 neurons^24^. The most prominent feature of a translating object is a leading moving edge followed by a trailing moving edge with an opposite contrast change. However, these moving edges can be confused by background moving edges. To increase the model’s selectivity, instead of directly calculating the correlation between the delayed leading edge and trailing edge signals as in the classical STMD model, we implemented an inhibition of inhibition mechanism, which proved particularly useful for selecting a translating object.

The inhibition of inhibition means that T2 and T3 are both inhibited by their own and each other’s signals (Fig. 2d). T2 and T3 first inhibit themselves through a self-inhibition branch, which receives inhibition from the delayed counterpart’s signal (denoted by 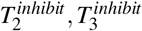). When a translating object is present, the leading edge’s signal is delayed to inhibit the inhibition branch of the trailing edge signal, resulting in a prominent signal. On the other hand, when no trailing edge follows, T2 and T3 signals are completely inhibited by themselves, preventing the model’s responses to single moving edges in the background. The outputs from the inhibition of inhibition mechanism are then compressed using a Tanh nonlinearity to form a small object map (SOM), representing the traces of the object’s moving edges:

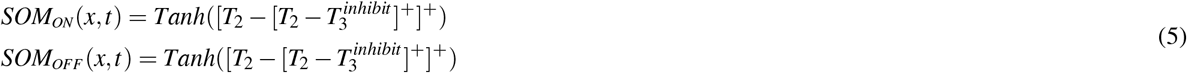

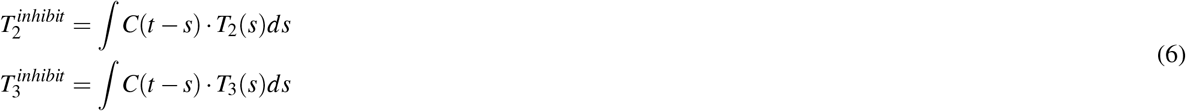

where *SOM*_*i*_ stands for small object map. The parameters *x* and *t* for *T*_2_ and *T* 3 are omitted for simplification. *C* is the temporal filter to delay *T*_2_ and *T*_3_ signals.

Note that T2 and T3 pathways have two *SOMs* for the ON/OFF moving target traces, respectively. A simple competition mechanism is adopted to select the maximum value between the two *SOMs*:

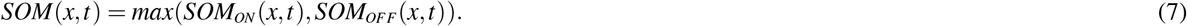

Upper in the pathways, T2 and T3 neurons receive presynaptic signals from fast-responding neurons Mi3, etc., in the Medulla. Thus, we model T2 and T3’s function by temporal filtering Medulla neuron outputs:

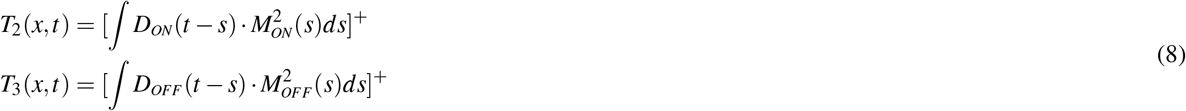

*D*_*ON/OFF*_ is the temporal filter ahead of T2 and T3 cells

#### The looming object detection branch

The looming detection branch is a key aspect of our paper (orange branch in Fig. 2a). We created a looming object map (LOM), marking potential looming objects based on inputs from the motion detection and/or feature detection pathways. This leads to three different looming detection models: one using only motion detection inputs, another with only edge feature detection inputs, and a third model with inputs from both pathways, called the combined-pathway model. In the combined-pathway model, we calculated the looming object map by multiplying the interneuron’s signals, which are the optic flow map and the edge feature map composed of spatial-temporally filtered T2/T3 signals. In the motion-pathway model, the LOM is the same as the optic flow map, while in the feature-pathway, it reduces to the edge feature map. The multiplication could be reinforcing and exclusive depending on whether the signals correlate. Hence, the unwanted signals from the background can be filtered out by the motion or the feature pathway, enhancing the model’s looming object detection capability in a dynamic background.

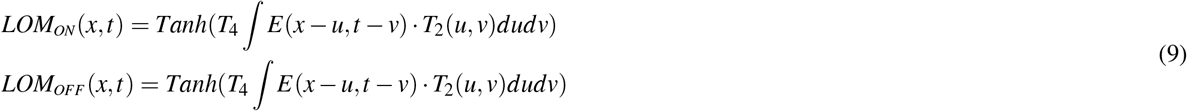

where *LOM*_*i*_ stands for looming object map. The two channels *LOM* are also compressed into one channel by competition:

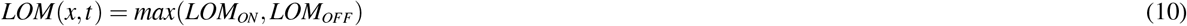

Unlike standard looming detection models that only give binary outputs (zero or one) to signal the presence of a looming object, we utilize two two-layer convolution networks to map the looming object map (LOM) into the size and speed of the looming object, respectively. There are two reasons for this approach:

1. An alarming model is limited in its predictive capabilities. Existing evidence suggests that individual neurons can encode information about looming size or speed, indicating that animals might perform more complex computations than just alarming.
2. It is relatively straightforward to design or train a network to achieve high accuracy as a binary classifier (zero or one). However, in such cases, it becomes challenging to demonstrate the additional benefits of combining motion and feature detection pathways for looming detection.

However, we must emphasize that apart from implementing four separate channels to account for the various object sizes, the convolution network was intentionally set simple with only a hidden layer to avoid taking over the roles of pattern recognition from the bioinspired models (Fig. 2e).

At last, all filters for the interneurons, including *A, B, C, D, E*, as well as the filters and CNNs in three separate branches, have free parameters to be trained. It’s essential to note that they are not initialized to function according to their names. Our objective is to demonstrate that by combining optimization tasks and an anatomy-constrained network, we can achieve a network that exhibits similar neural dynamics to their biological counterparts, specifically, T2, T3, T4, and T5 neurons. We also aim to show that our network can converge to a biological solution by hypothetically multiplying the motion and feature detection signals, validating our approach to model the coordination between the two pathways.

#### Dataset

We selected 241 high-quality panoramic high dynamic range images from the original 421 images^41^ to use as backgrounds in our video sequences. To simulate the ego motion, we assumed that the insect remains stationary, but the entire background moves as if a moving insect is turning. The background’s movement speed follows a stochastic sequence sampled from a Gaussian distribution with a mean of 0 and a standard deviation of 90°/s. Additionally, we manually introduced moving objects in the foreground, including both translating and looming objects. The largest object size covers a visual field of 120°. To save computational resources, the high-resolution images were down-sampled using spatial filters to match the resolution of a fly compound eye. Since we focused on 1D scenarios, we stimulated the model with the movement of a line of pixels. This motion lasted for 2 seconds and was sampled at 100Hz, resulting in a 2D spatial-temporal pattern with 72*200 data points (Fig. **??**a right).

To mimic the wide range of contrast distributions seen in natural objects, the contrast distribution of the added objects is more diverse than that of the background (Extended Data Fig. 2). Then the Weber contrast of the normalized background scene is slightly biased with a zero mean and a standard variation^42^. The variation of the object’s Weber contrast is about three times larger.

The Weber contrast is defined as:

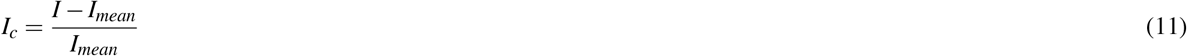

#### Optimization and evaluation

We trained our model in a supervised way to optimize a multi-objective loss function that considers the background moving direction, the looming object size, the looming object speed and the translating object speed. To ensure unbiased training, we sample the same number of datasets with looming objects and translating objects. We used mean square error (MSE) as the training loss in optimization and R-square to evaluate models’ performances. The loss function is defined as:

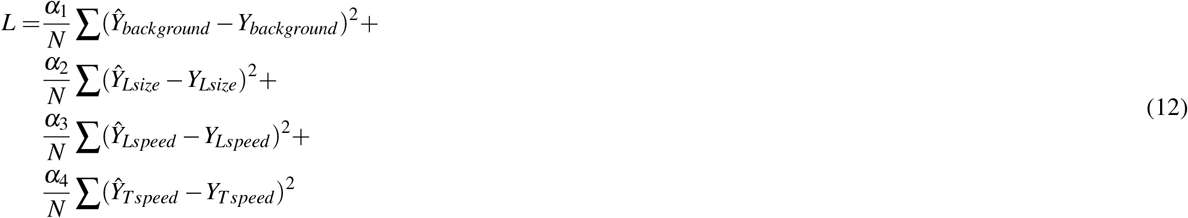

where *α*_*j*_, *j ∈* 1, 2, 3, 4 are the weights for the four tasks. In our training process, they were set as 100, 1, 1, and 1. *Y*_*background*_ is the background motion direction formulated by values between -1 and 1. *Y*_*Lsize*_ and *Y*_*Lspeed*_ are the size and speed of the looming object. *Y*_*Tspeed*_ is the speed of the translating object. And the hat above each *Y* stands for the model estimations.

The evaluation R-square is defined by:

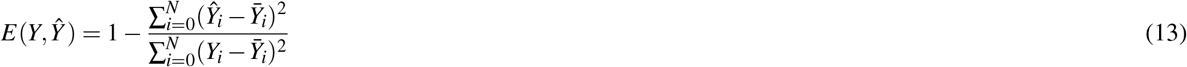

where *E* is the evaluation for the looming size estimation. *Y*_*i*_ is the label and *Ŷ*_*i*_ is the model estimation. 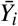is the mean value of *Y*_*i*_. The *E* value is up-bounded by 1, and the larger its value, the better the model in estimating the labelled values.

## Data Availability

The natural image database used in this study has DOI https://pub.uni-bielefeld.de/record/2689637 and is available at https://pub.uni-bielefeld.de/rc/2689637/2693616.

## Code Availability

All code for experiments and figures generation in Python is available at xxxxxxxx for the purposes of reproducing and extending the analysis.

## Acknowledgements

We thank Prof. Marion Sillies, Prof. Shigang Yue, Dr. Yu Zhou, and Dr. Jian Liu for valuable feedback on the initial drafts of the manuscripts. Project supported by the Young Scientists Fund of the National Natural Science Foundation of China (Grant No. 12001111), Fund from the National Natural Science Foundation of China (Grant No. 11925103), Shanghai Municipal Science and Technology Major Project (No.2018SHZDZX01), ZJ Lab, and Shanghai Center for Brain Science and Brain-Inspired Technology, the 111 Project (No.B18015), and the 2021 STCSM (Grant No. 2021SHZDZX0103)

## Author contributions statement

B.G. and Z.S. designed the study. B.G. programmed the model and ran the simulations, with supervision from Z.S.; Z.S. wrote the paper with an initial draft from B.G. and editing from everybody; B.G. and Z.S. analysed the results and designed the figures. JF.F advised the project.

## Competing interests

The authors declare no competing interest.

**Extended Data Fig. 1.**
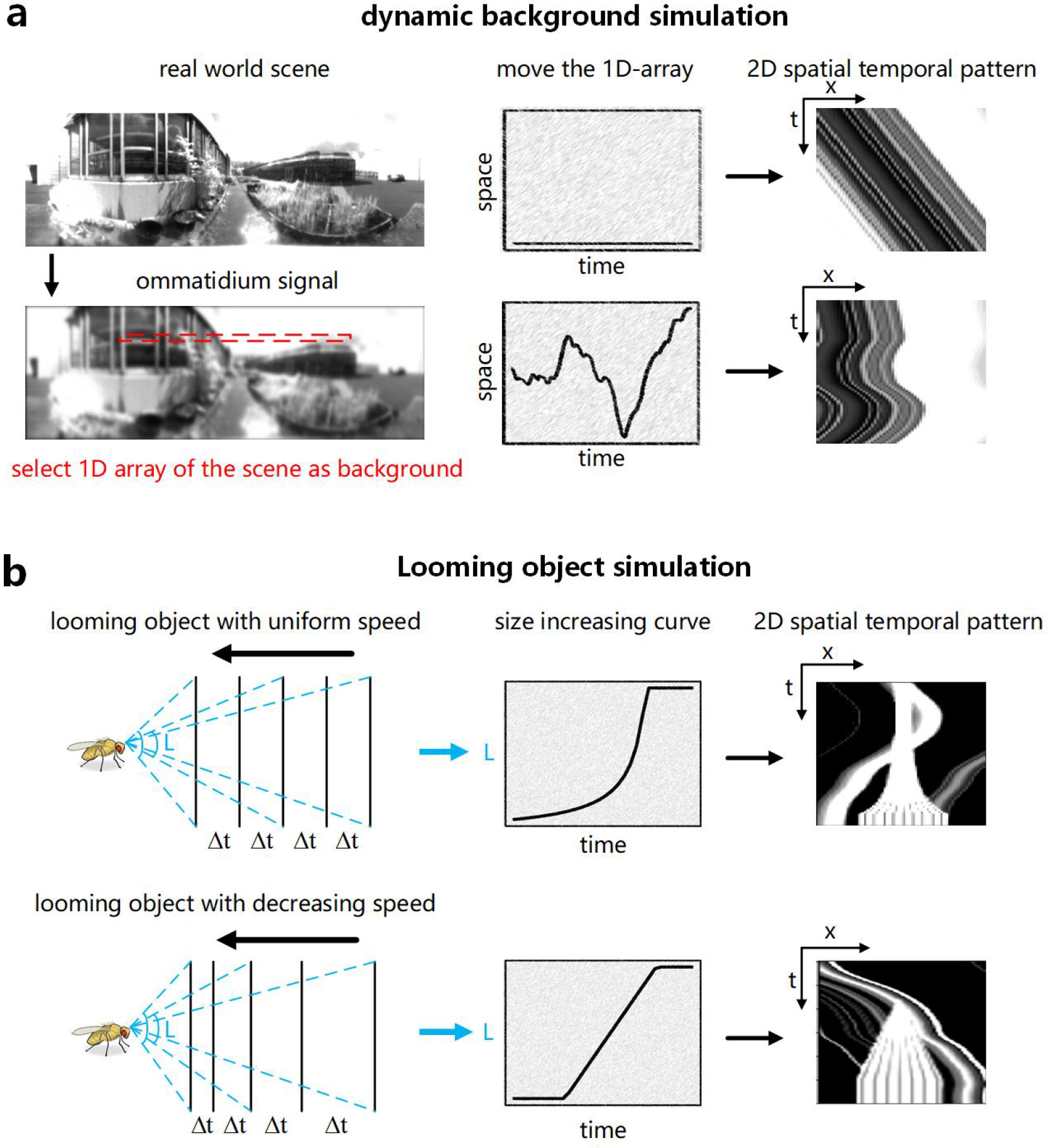
The procedure to generate datasets. (a) A selected panoramic image of a natural scene; (b) A 1-D array of signals from the low-pass filtered image; (c) A spatiotemporal signal of the 1-D image that moves at a given speed; (d) The spatiotemporal signals when an ON/OFF contrasted looming object or a translating object is added to (c); (e) Weber-contrast distribution of the background scenes and the manually added object.

**Extended Data Fig. 2.**
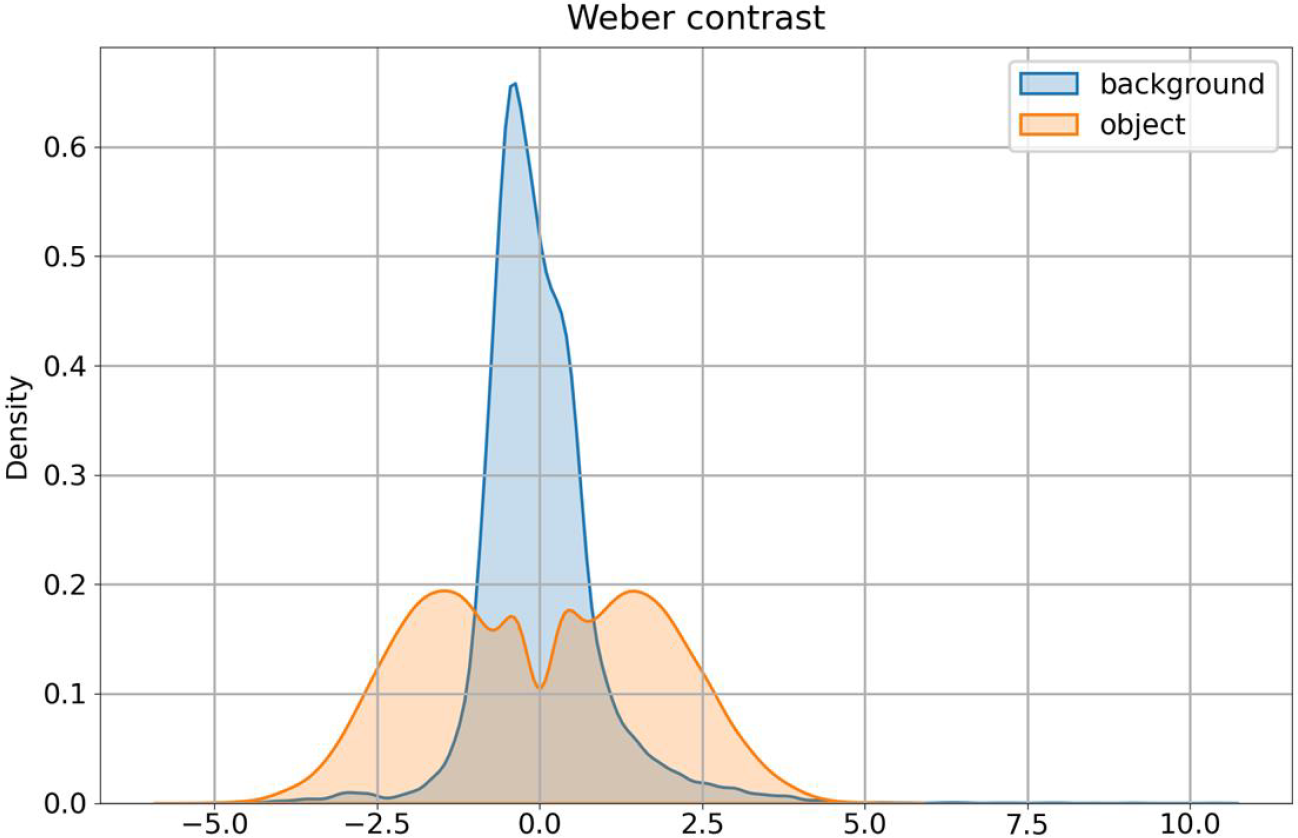
Weber contrast distribution of the background scenes and manually added object.

**Extended Data Fig. 3.**
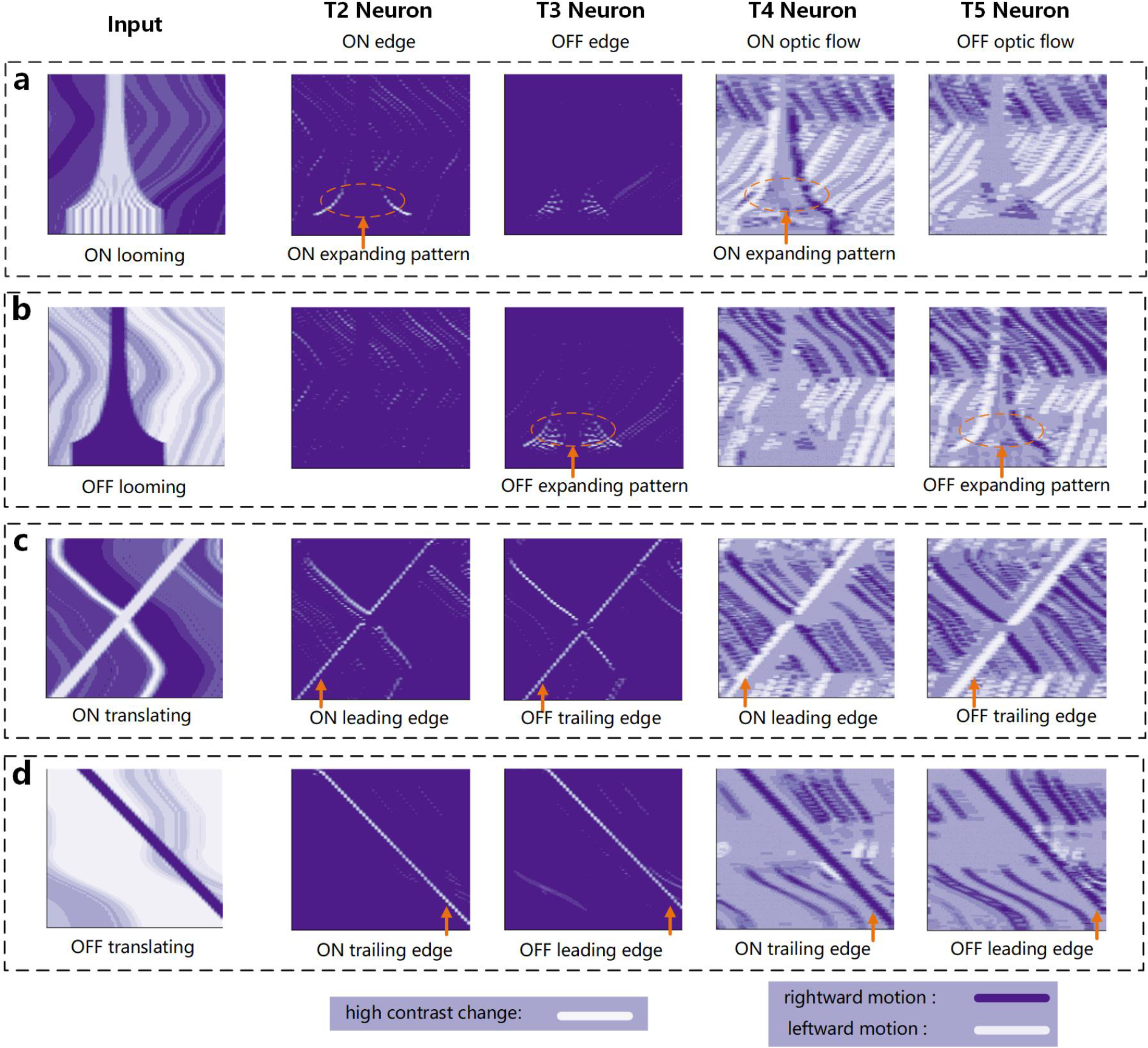
The interneurons function as expected, similar to their biological counterparts. T2 and T3 separate into ON/OFF pathways like T4 and T5 cells. T2 and T4 cells respond strongly to ON looming objects, while T3 and T5 cells are selective for OFF looming objects. T2 and T3 focus on local high-contrast change signals with less background noise, while T4 and T5 extract local motion signals and retain motion direction opponency. (a) Interneuron response map for an ON looming object with a sine-wave pattern, T2 and T4 neuron responses show a clear expanding pattern. T2’s responses are less noisy but appear towards the end of the looming phase, while T4’s responses are more noisy but exhibit larger looming responses. (b) Interneuron response map for an OFF looming object with a sine-wave pattern. (c) Interneuron response map for an ON translating object. (d) Interneuron response map for an OFF translating object.

**Extended Data Fig. 4.**
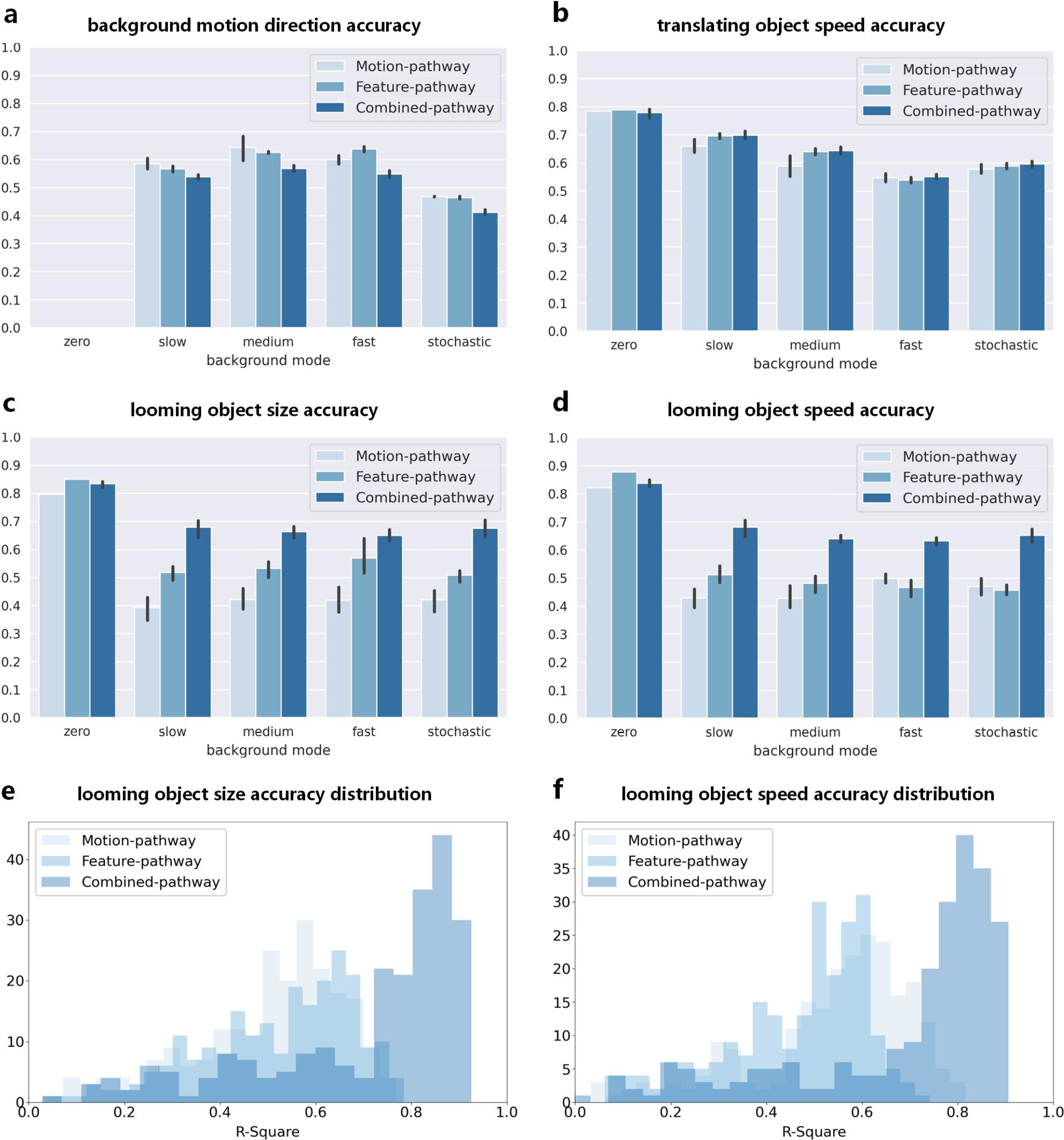
Comparisons of three looming detection model performances in static or dynamic backgrounds with linear expanding objects, moving stochastically or with uniform speeds at different ratios of the linearly expanding looming speed. All three models’ performances decline significantly when the background is moving. However, the combined-pathway model enhances looming detection performances in all tested dynamic conditions. (a b) The three models show similar performance for estimating background motion direction and translating object speed, indicating that the complementary mechanism does not negatively affect these two branches’ functions. (c d) The complementary model improves the accuracy of estimating looming object size and speed in all tested dynamic conditions but not in static backgrounds. (e f) Performance distributions for looming size and speed estimation for the three models tested with stochastic moving backgrounds.

## Notes

### Competing Interest Statement

The authors have declared no competing interest.

